# Mouse mutants in schizophrenia risk genes *GRIN2A* and *AKAP11* show EEG abnormalities in common with schizophrenia patients

**DOI:** 10.1101/2022.04.05.487037

**Authors:** Linnea E. Herzog, Lei Wang, Eunah Yu, Soonwook Choi, Zohreh Farsi, Bryan Song, Jen Q. Pan, Morgan Sheng

## Abstract

**BACKGROUND:** Schizophrenia is a heterogeneous psychiatric disorder with a strong genetic basis, whose etiology and pathophysiology remain poorly understood. Exome sequencing studies have uncovered rare, loss-of-function variants that greatly increase risk of schizophrenia [1], including loss-of-function mutations in *GRIN2A* (aka *GluN2A* or *NR2A*, encoding the NMDA receptor subunit 2A) and *AKAP11* (A-Kinase Anchoring Protein 11). *AKAP11* and *GRIN2A* mutations are also associated with bipolar disorder [2], and epilepsy and developmental delay/intellectual disorder [1, 3, 4], respectively. Accessible in both humans and rodents, electroencephalogram (EEG) recordings offer a window into brain activity and display abnormal features in schizophrenia patients. Does loss of *Grin2a* or *Akap11* in mice also result in EEG abnormalities?

**METHODS:** We monitored EEG in heterozygous and homozygous knockout *Grin2a* and *Akap11* mutant mice compared with their wild-type littermates, at 3- and 6-months of age, across the sleep/wake cycle and during auditory stimulation protocols.

**RESULTS:** *Grin2a* and *Akap11* mutants exhibited increased resting gamma power, attenuated 40-50 Hz auditory steady-state responses (ASSR), and reduced responses to unexpected auditory stimuli during mismatch negativity (MMN) tests. Sleep spindle density was reduced in a gene dose-dependent manner in *Akap11* mutants, whereas *Grin2a* mutants showed increased sleep spindle density.

**CONCLUSIONS:** The EEG phenotypes of *Grin2a* and *Akap11* mutant mice show a variety of abnormal features that overlap considerably with human schizophrenia patients, reflecting systems-level changes caused by *Grin2a* and *Akap11* deficiency. These neurophysiologic findings further substantiate *Grin2a* and *Akap11* mutants as genetic models of schizophrenia and identify potential biomarkers for stratification of schizophrenia patients.

## INTRODUCTION

A severe psychiatric disorder affecting approximately 0.5% of the global population, schizophrenia is characterized by hallucinations, delusions, disorganized thoughts and behavior, social withdrawal, reduced emotional expression, and cognitive deficits. While genome-wide association studies (GWAS) have identified many genetic loci associated with schizophrenia, it is often difficult to identify the causal gene or to interpret the biological effect of the GWAS common variants, which are mainly non-coding, challenging to fine-map, and associated with only small increases in disease risk (odds ratio typically around 1.1) [5-7].

Compared with GWAS, the rare copy number variants that are associated with schizophrenia confer much higher risk but affect large regions of the genome, making it difficult to identify the relevant pathogenic gene(s) [7, 8]. As a result, the field has been hampered by lack of specific genetic models that can be brought to bear on the mechanisms of schizophrenia.

Recently, the discovery of schizophrenia risk genes has been enhanced by large-scale exome or genome sequencing of tens of thousands of cases versus controls. Such studies have the power to uncover rare loss-of-function coding variants (such as protein truncating variants) that have a large impact on schizophrenia risk [1, 9, 10]. In one of the largest sequencing studies to date of 24,248 cases and 97,322 controls, the Schizophrenia Exome Sequencing Meta-analysis Consortium (SCHEMA) has identified multiple rare loss-of-function genetic variants at exome-wide level of significance that confer substantial disease risk (“SCHEMA genes”; odds ratios in the range of 4-50) [1]. Because these rare variants are often protein-truncating, thus presumably loss-of-function null mutations [1, 11], these disease-causing mutations can be easily modeled by genetic disruption (‘knockout’) in animals such as the mouse. Moreover, these null mutations can be studied in homozygous as well as heterozygous states, the latter being more relevant to the human disease, where only one of the alleles is disrupted in patients. By systematically analyzing the phenotypes of mouse lines bearing loss-of-function mutations in SCHEMA genes (SCHEMA mouse mutants), we hope to discover convergent molecular and neurobiological mechanisms and identify potential biomarker signatures caused by these relatively highly penetrant mutations. Insights derived from these genetic animal models of schizophrenia should further our understanding of the biology of schizophrenia and could aid our efforts to develop more effective treatments [12]. In this study, we characterize by chronic electroencephalogram (EEG) the changes in brain activity caused by mutations in two SCHEMA genes, *GRIN2A* and *AKAP11*.

*GRIN2A* (glutamate ionotropic receptor NMDA type subunit 2A) is a particularly compelling SCHEMA gene for further study because it is also a significant GWAS hit for schizophrenia [5]. *GRIN2A* encodes the 2A subunit of the N-methyl-D-aspartate (NMDA) receptor (also known as GluN2A or NR2A), a protein that is highly expressed in neurons in the brain and localized at postsynaptic sites of glutamatergic synapses [13, 14]. Reduced NMDA receptor function has long been proposed as a pathophysiologic mechanism underlying schizophrenia, in part because NMDA receptor antagonists at low concentrations can induce psychosis-like symptoms in humans [15]. Interestingly, *GRIN2A* is expressed later in brain development than the genes encoding the GRIN1 and GRIN2B subunits of the NMDA receptor; expression of *GRIN2A* starts postnatally and rises through juvenile and adolescent stages in humans and rodents, inviting comparison with the typical onset of schizophrenia in adolescence and early adulthood [16-18].

*AKAP11* (A-Kinase Anchoring Protein 11) has been identified as a rare-variant large-effect risk gene for schizophrenia in the SCHEMA study [1]. In addition, *AKAP11* is a bipolar disorder risk gene, recently identified as the top hit from Bipolar Exome (BipEx) sequencing studies of 13,933 bipolar cases and 14,422 controls [2]. Thus, *AKAP11* is a shared risk gene for schizophrenia and bipolar disorder, underscoring the genetic overlap between these two disorders on the psychosis spectrum. *AKAP11* (formerly known as *AKAP220*) encodes an A-kinase anchoring protein that is expressed broadly in the body, including in neurons in the brain. Biochemically, AKAP11 interacts with protein kinase A (PKA), a protein that is involved in a variety of biological processes including neuronal plasticity, and with glycogen synthase kinase 3 beta (GSK3β), one of the targets of lithium, a mainstay therapy for bipolar disorder [19-21]. However, AKAP11’s function in the brain and its biological role in psychiatric disease are uncharacterized.

Here we investigate how *GRIN2A* and *AKAP11* loss-of-function affect brain activity by chronic monitoring of EEG in mutant mice deficient in these genes. EEG provides a non-invasive, high temporal-resolution, systems-level readout of neural activity *in vivo*, a neurophysiologic assay that is clinically translatable to human patients. The present work provides an in-depth characterization of EEG phenotypes across different behavioral states (sleep, wake, auditory stimulation) and ages (3 and 6 months) in *Grin2a* and *Akap11* and heterozygous (Het) and homozygous knockout (KO) mice, as compared with their wild-type littermates (WT). We found that *Grin2a* and *Akap11* mutant mice possess a number of EEG features shared with human schizophrenia and bipolar disorder patients, including elevated gamma oscillations at rest [22, 23], attenuated auditory steady-state responses (ASSR) at 40-50 Hz [24, 25], and changes in sleep spindle density [26, 27]. These mouse EEG phenotypes reveal systems-level abnormalities caused by (even heterozygous) mutations in *Grin2a* and *Akap11*, and provide further justification, beyond their genetic validity, that these mouse mutants can serve as useful animal models of schizophrenia/bipolar disorder.

## MATERIALS AND METHODS

### Animals

All experiments were approved by the Broad Institute IACUC (Institutional Animal Care and Use Committee) and conducted in accordance with the NIH Guide for the Care and Use of Laboratory Animals. Mice were housed at AAALAC-approved facilities on a 12-hour light/dark cycle, with food and water available *ad libitum. Grin2a* (B6;129S-Grin2a<tm1Nak>; Riken BioResource Center, RBRC02256) and *Akap11* (B6.Cg-Akap11[tm1.2Jsco/J]; Jackson Laboratory, #028922) knockout mice were originally generated as described [28, 29] and maintained in-house by breeding with C57BL/6J wild-type mice (Jackson Laboratory, #000664). The resulting heterozygous breeding pairs were used to generate adult male and female *Grin2a* and *Akap11*^*-/-*^ (knockout, KO) and ^+/-^ (heterozygous, Het) mice and their wild-type (^+/+^, WT) littermates, which were used for all experiments. Open field behavior and auditory EEG experiments were conducted during the light phase of the daily cycle. All experiments and analyses were conducted by investigators who were blinded to the mouse genotype.

### Open field testing

11-to 14-week-old *Grin2a* and *Akap11* mutant mice and their WT littermates (*n*=10-13 animals/group) were monitored using the SuperFlex Open Field system (Omnitech Electronics, Inc., 40cm x 40cm x 40cm) for 60 minutes. The animals’ position was captured in real-time using Fusion system software (Omnitech Electronics, Inc., Columbus, OH). Total distance and the ratio of center/margin distance for each group were analyzed using one-way ANOVAs with post hoc Tukey tests.

### EEG implantation surgery

6-to 12-week-old mice (*n=*11-12 mice/group) were deeply anesthetized with isoflurane. A prefabricated EEG/EMG headmount (#8201-SS, Pinnacle Technologies) was secured to the skull with four 0.10’’ intracranial electrode screws (#8403, Pinnacle Technologies) at the following stereotactic coordinates: frontal recording electrode (+1.5 AP, 1.5 ML to Bregma), parietal recording electrode (−2 AP, 1.5 ML to Bregma), ground and reference electrodes (bilaterally -1 AP, 2 ML to Lambda). The electromyogram (EMG) electrodes were placed bilaterally in the nuchal muscles. Electrodes were soldered to the EEG/EMG headmount and dental acrylic was used to secure the connections. Animals were given at least one week of post-operative recovery before EEG recording. This manuscript reports results from the frontal EEG electrode only.

### EEG recording

Following recovery from EEG implantation, mice were tethered to the Pinnacle recording system, with at least 3 hours of habituation before testing. Experiments were conducted over a period of 4-5 days. EEG/EMG signals were recorded in freely moving mice across 24 hours of sleep/wake, followed by ASSR and mismatch negativity (MMN) testing (see Supplementary Information for details).

Animals remained tethered to the Pinnacle system throughout the testing period with *ad libitum* access to food and water. All signals were digitized at a sampling rate of 1000 Hz, filtered (1– 100 Hz bandpass for EEG; 10–1 kHz bandpass for EMG), and acquired using the Sirenia Acquisition program (Pinnacle Technologies). EEG recordings took place at two timepoints, roughly corresponding to 3- and 6-months of age (mouse age at time of Recording 1: 8-17 weeks, Recording 2: 22-33 weeks). Mice were returned to their home cage between recording sessions.

### EEG analysis

Sleep/wake power, ASSR, and MMN were analyzed as described in the Supplementary Information.

### Code availability

LUNA software (http://zzz.bwh.harvard.edu/luna) was used for sleep/wake EEG analysis. All other analyses were conducted using custom Python and MATLAB (MathWorks, Natick, MA, RRID: SCR_001622) scripts, which the authors will provide upon request.

## RESULTS

### Grin2a and Akap11 mutants exhibit increased and decreased locomotor activity, respectively, in the open field test

The open-field test is frequently used to measure general locomotor activity and infer anxiety-like behavior in rodents [30, 31]. At 3 months of age, *Grin2a* KO mice showed increased locomotor activity in the open field (∼25% increase from WT; *p*=0.033), while *Grin2a*^+/-^ mice exhibited similar activity levels as WT littermates (Figure 1A, C, E). In contrast, *Akap11* Het (*p*=0.0337) and KO (*p*=7.49e-07) showed reduced locomotor activity ∼by 25% and 50%, respectively, compared to their WT littermates (Figure 1B, D, G), consistent with previous reports [29, 32]. *Grin2a* KO (but not Het) mice had reduced ratios of center/margin distance traveled (by ∼30%) relative to WT animals (*p*=0.0438), suggesting anxiety-like behavior (Figure 1F). *Akap11* Het and KO had similar center/margin ratios as WT littermates (Figure 1H).

**Figure 1.**
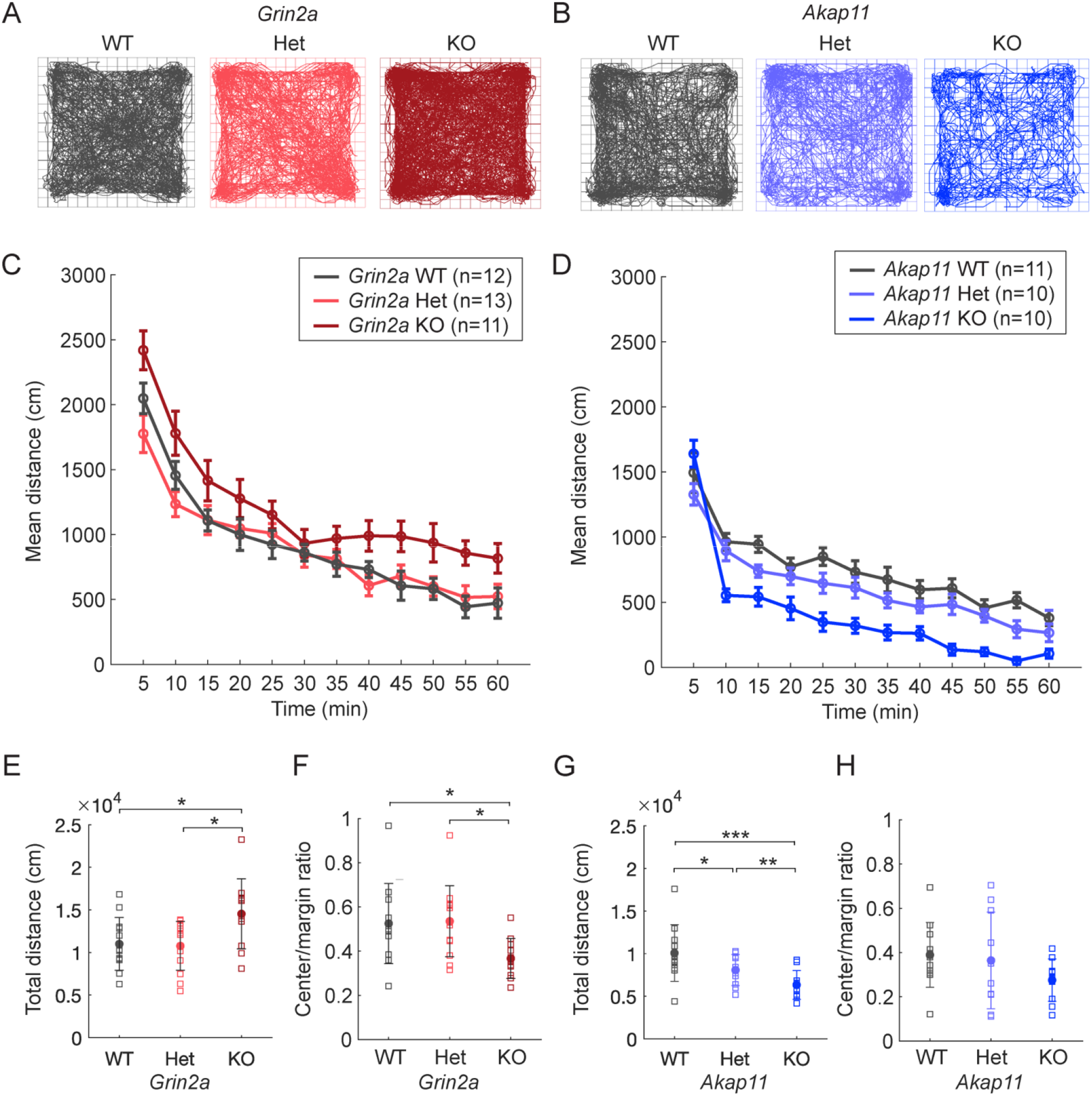
Locomotor behavior of *Grin2a* and *Akap11* mutant mice in open field test. **(A-B)** Representative traces of mouse movements (60-minute test). **(C-D)** Mean distance traveled, binned into 5-minute segments. **(E-H)** Total distance traveled and ratio of center/margin distance in *Grin2a* (left) and *Akap11* (right) mutants in comparison to WT littermates. Error bars denote mean +/- standard error; \**p<*0.05, \*\**p<*0.01, \*\*\**p*<0.001.

### *Akap11*^*-/-*^ mice exhibit NREM sleep deficits

Patients with schizophrenia often show sleep abnormalities, including reduced sleep, increased sleep, delayed sleep onset, or fragmented sleep, relative to healthy subjects [33, 34]; altered sleep patterns (e.g. more sleep in depressive phase, less sleep in manic phase) are also found in bipolar disorder [35]. To assess sleep differences in *Grin2a* and *Akap11* mutant mice, we recorded EEG/EMG signals across 24 hours and used a feature-based model (see Supplementary Information) to classify periods of NREM, REM, and wake. Sleep analyses were conducted during the light cycle when mice are predominantly asleep. *Grin2a* heterozygous and homozygous mutant mice exhibited similar sleep patterns as their WT littermates (Figure 2A-B). *Akap11*^-/-^ mice exhibited reduced (∼10%) NREM sleep relative to WT littermates at 3 months (*p*=0.0052) and 6 months of age (*p*=0.0106), whereas *Akap11*^*+/-*^ animals were not significantly different than WT (Figure 2C-D). At 6-months of age, *Akap11*^-/-^ mice also showed delayed sleep onset at the start of the light cycle, relative to WT littermates (*p*=0.0207, Figure 2H). Neither *Grin2a* or *Akap11* mutants (heterozygous or homozygous) displayed abnormal sleep fragmentation (Figure S1), although NREM bout length was reduced in *Akap11* KO (*p*=0.0478) compared with WT littermates (Figure S1G).

**Figure 2.**
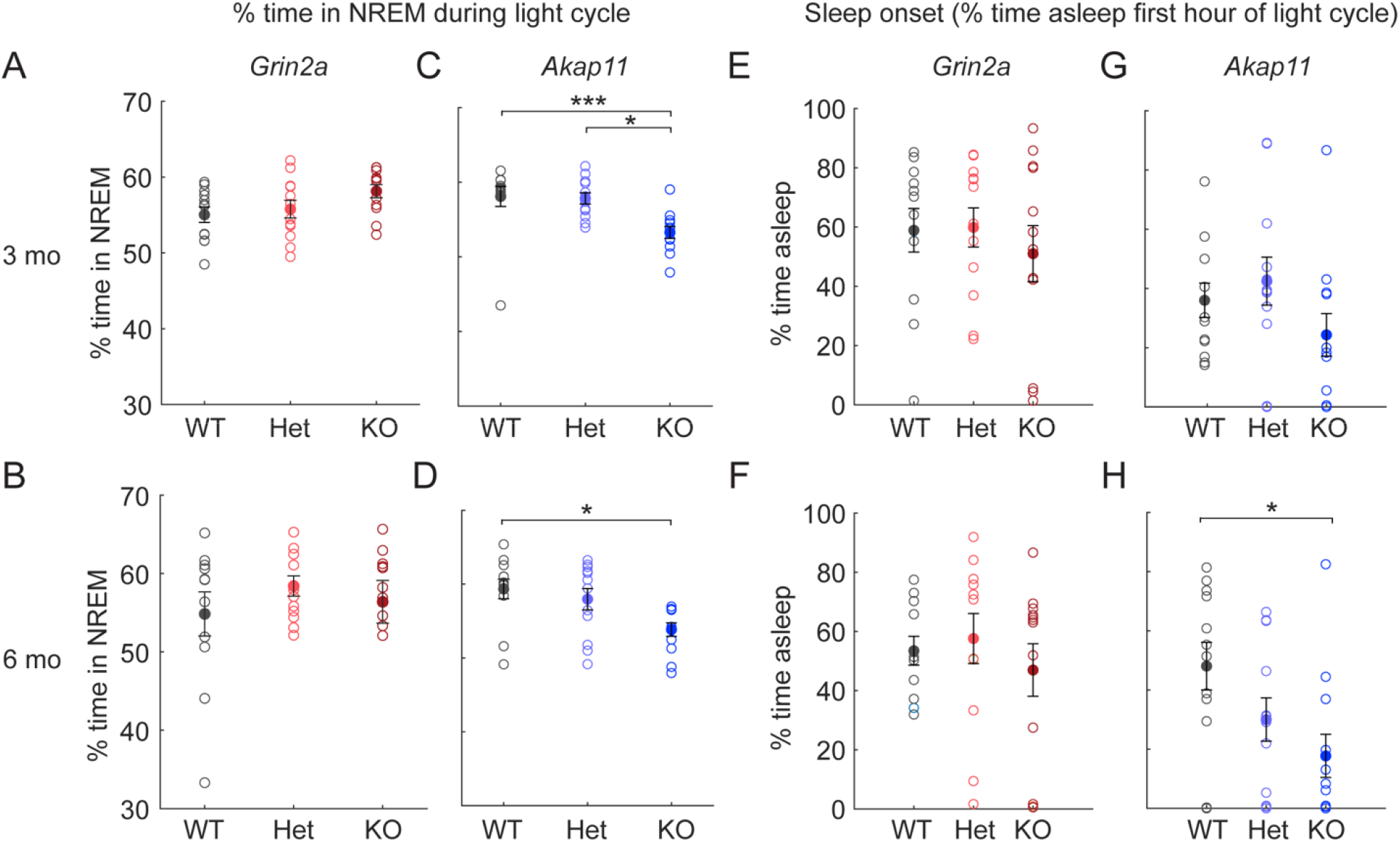
NREM sleep and sleep onset time of *Grin2a* and *Akap11* mutant mice. **(A-D)** % of time spent in NREM sleep during the light cycle in 3- and 6-month *Grin2a* and *Akap11* mutants. **(E-H)** Sleep onset, as quantified by the % of time spent asleep in the first hour of the light cycle. Error bars denote mean +/- standard error; *p<*0.05, \*\**p<*0.01, \*\*\**p*<0.001; *n=*11-12 mice/group.

### Elevated resting gamma power in *Grin2a* and *Akap11* mutant mice

Many patients with schizophrenia exhibit altered EEG oscillatory power at rest, especially increased gamma oscillations during sleep and quiet wake [36, 37]. Focusing our analysis on NREM sleep, where EEG signals are less prone to movement-related variation, we measured absolute power of brain oscillations in *Grin2a* and *Akap11* mutants versus their WT littermates for each frequency band (slow: 0.5-1 Hz, delta: 1-4 Hz, alpha: 8-12 Hz, sigma: 12-15 Hz, beta: 15-30 Hz, gamma: 30-50 Hz). *Grin2a*^+/-^ and ^-/-^ mice exhibited gene dose-dependent increases in gamma oscillations (Figure 3A-B; Figure S2E, J): a 10% and ∼20% increase in absolute gamma power compared with WT for heterozygous and homozygous mutants, respectively (Het, 3 months: *p*=0.0187; Het, 6 months: *p*=0.0385; KO, 3 months: *p*=1.07e-08; KO, 6 months: *p*=1.15e-04). Resting gamma power was also increased by ∼10% in *Akap11*^*-/-*^ mice at 3 months (*p*=1.80e-04) and 6 months (*p*=0.0112) of age, but unlike *Grin2a* heterozygotes, *Akap11* heterozygous mutants were not significantly different from WT (Figure 3C-D; Figure S2N, P). Besides NREM, elevated gamma power was also found during REM sleep (Figure S2; Figure S3; Figure S4C, F, J, L) and quiet wake (Figure S5) in *Grin2a* and *Akap11* homozygous mutants. *Grin2a* (but not *Akap11*) heterozygous mutants exhibited an intermediate phenotype of increased gamma oscillations during REM sleep, although this result was not statistically significant (3 months: *p=*0.190; 6 months: *p=*0.111) or quiet wake (3 months: *p*=0.196; 6 months: *p=*0.872).

**Figure 3.**
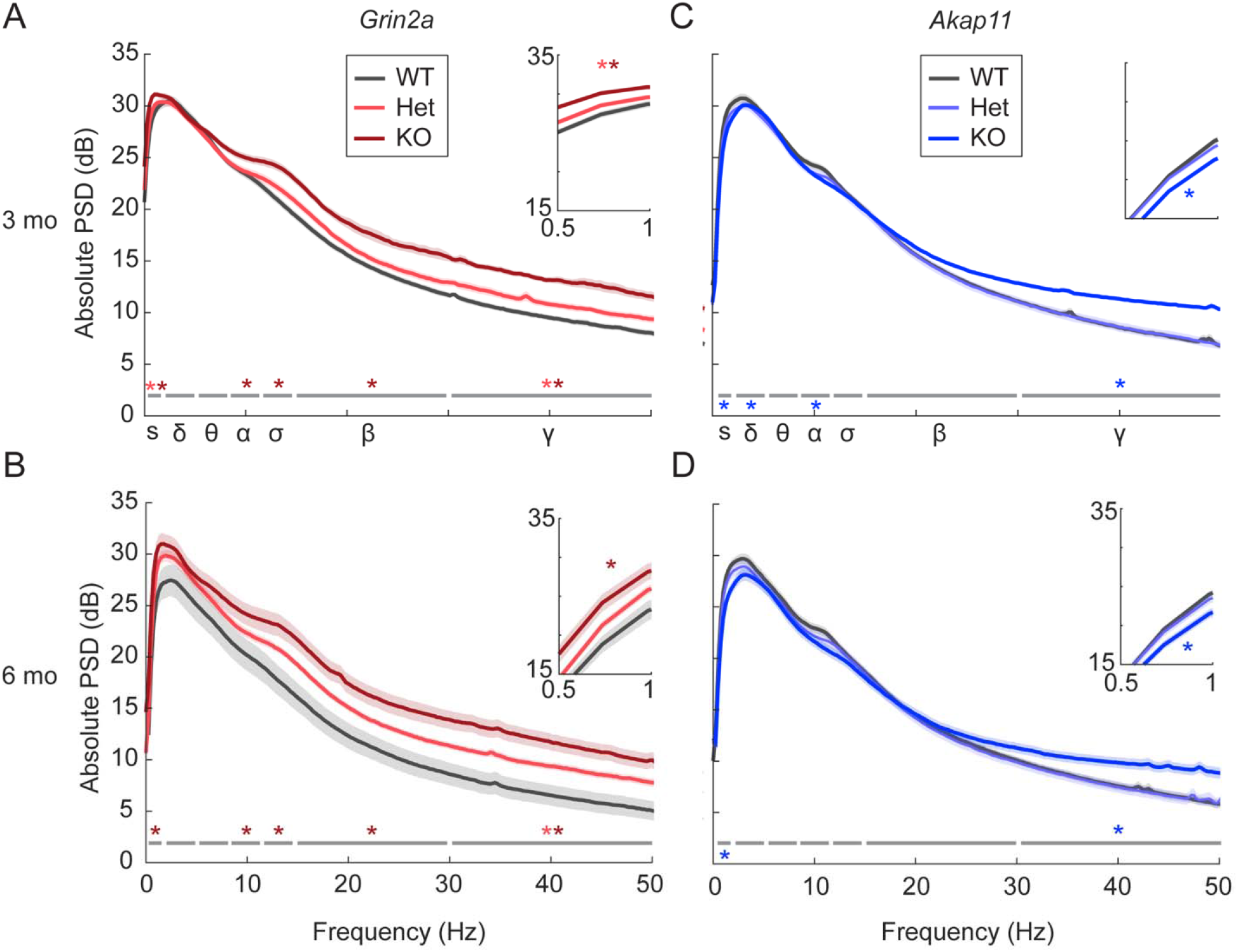
Power spectral analysis of brain oscillations in *Grin2a* and *Akap11* mutant mice during NREM sleep. **(A-D)** Absolute NREM power spectra for 3- and 6-month *Grin2a* and *Akap11* mutants during the light cycle. Light and dark red/blue asterisks indicate oscillations for which ^+/-^ and ^-/-^ mice, respectively, differed significantly from WT littermates (*p*<0.05) in each of the frequency bands: slow (s), 0.5-1 Hz; delta (δ), 1-4 Hz; theta (θ), 4-8 Hz; alpha (α), 8-12 Hz; sigma (δ), 12-15 Hz; beta (β), 15-30 Hz; gamma (γ), 30-50 Hz; with asterisks above or below the gray line indicating significant increases or decreases in power, respectively. Shading indicates mean +/- standard error. Insets show magnified view of the graph in the slow oscillation range (0.5-1 Hz). *n=*11-12 mice/group.

We found changes in oscillation power outside of gamma frequencies. The EEGs of *Grin2a* KO exhibited increased power broadly across multiple frequency bands, including slow (3 months: *p=*8.12e-08; 6 months: *p=*0.0023), alpha (3 months: *p*=0.0252; 6 months: *p*=0.0357), sigma (3 months: *p=*4.00e-05; 6 months: *p=*0.0030), and beta (3 months: *p*=2.71e-06; 6 months: *p*=7.76e-04) (Figure 3A-B; Figure S2A-J). *Grin2a* Hets also exhibited increased power of slow oscillations at 3 months (*p*=0.0259). Unlike *Grin2a* KO mice, *Akap11*^*-/-*^ animals had reduced power of slow oscillations (Figure 3C-D; S2K), by ∼15% compared to WT (3 months: *p*=2.85e-05; 6 months: *p*=0.0059), as well as reduced delta (*p=*0.0191) and alpha (*p=*0.0375) oscillations at 3 months (Figure 3C-D; S2L-M). Heterozygous *Akap11* mutants showed no significant changes in oscillation power for any frequency band.

### *Grin2a* and *Akap11* mutants exhibit gene dose-dependent increases and decreases in sleep spindles, respectively

Sleep spindles, an EEG feature possibly linked to overnight memory consolidation, are commonly reduced in schizophrenia patients [38, 39]. Measuring sleep spindle density during NREM sleep in the light cycle, we found striking gene dose-dependent increases in sleep spindle density in *Grin2a* mutants, especially at 6 months (Figure 4A-B; ∼10-40% increase in heterozygous, and 25-50% increase in homozygous mutants). This increase in spindle density occurred across a range of spindle frequencies from 9 Hz to 15 Hz (3 months, KO, 15 Hz: *p*=0.0234; 6 months, Het, 13 Hz: *p*=0.0340; 6 months, Het, 15 Hz: *p*=0.0192; 6 months, KO, 9 Hz: *p=*0.0188; 6 months, KO, 13 Hz: *p=*0.0044; 6 months, KO, 15 Hz: *p*=0.0016). In contrast, *Akap11* mutants showed the opposite phenotype of decreased spindles (Figure 4C-D), also in a gene dose-dependent manner (heterozygotes ∼25% reduction and homozygous KO ∼50-75% reduction, compared with WT littermates) (3 months, Het, 9 Hz: *p*=0.0029, 3 months, Het, 11 Hz: *p=*0.0193; 3 months, KO, 9 Hz: *p*=5.53e-09; 3 months, KO, 11 Hz: *p*=6.58e-11; 3 months, KO, 13 Hz: *p*=2.27e-06; 3 months, KO, 15 Hz: *p*=2.61e-04; 6 months, Het, 9 Hz: *p*=0.0088; 6 months, Het, 11 Hz: *p=*0.0106; 6 months, KO, 9 Hz: *p*=1.03e-08; 6 months, KO, 11 Hz: *p*=1.39e-11; 6 months, KO, 13 Hz: *p*=1.00e-06; 6 months, KO, 15 Hz: *p*=1.52e-04).

**Figure 4.**
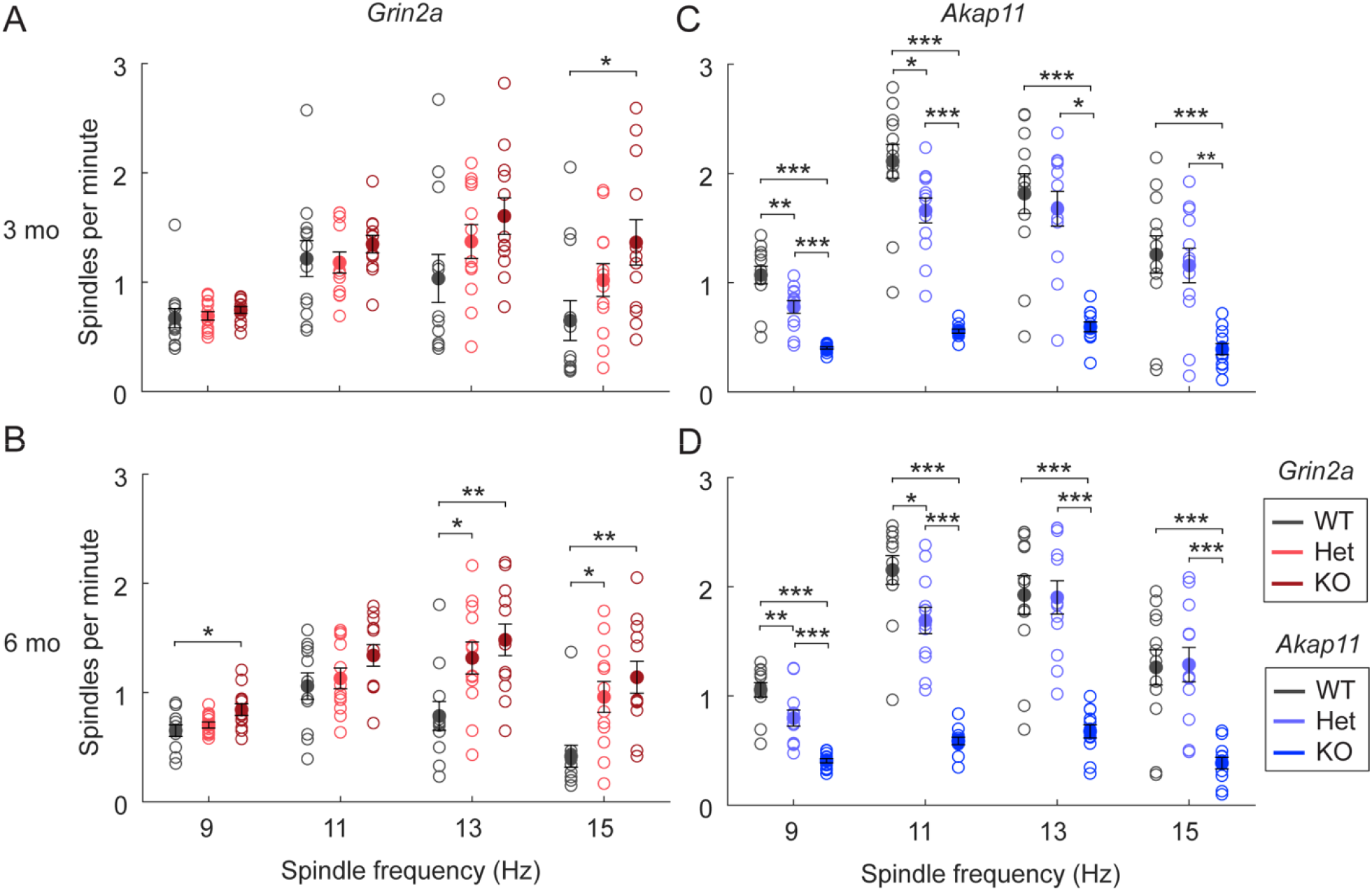
Sleep spindle density in *Grin2a* and *Akap11* mutants. **(A-D)** Density for 9 Hz, 11 Hz, 13 Hz and 15 Hz spindles during NREM sleep during the light cycle. Error bars show mean +/- standard error; *p<*0.05, \*\**p<*0.01, \*\*\**p*<0.001; *n=*11-12 mice/group.

### *Grin2a* and *Akap11* mutants exhibit attenuated ASSR at gamma frequencies

ASSR is a highly translatable sensory processing assay that measures cortical entrainment by comparing the evoked response to click trains of different frequencies relative to pre-stimulus activity [40]. ASSR is typically reduced for 40-50 Hz (gamma frequency) stimuli in schizophrenia and bipolar disorder patients relative to healthy controls, signifying impaired entrainment of cortical gamma rhythms [22, 25, 41, 42]. ASSR was reduced in *Grin2a* mouse mutants, particularly in the 40-50 Hz frequency range, and more prominently at 6 months of age (Figure 5A-B). Relative to WT, homozygous (*p*=0.0467) and heterozygous (*p*=0.0214) *Grin2a* mutants showed a ∼30% reduction in 50 Hz ASSR at 6 months. There was a trend (*p*=0.0521) towards decreased 40 Hz ASSR for *Akap11* KO at 3 months (Figure 5C), but otherwise, there were no consistent differences between *Akap11* mutants and WT (Figure 5C-D).

**Figure 5.**
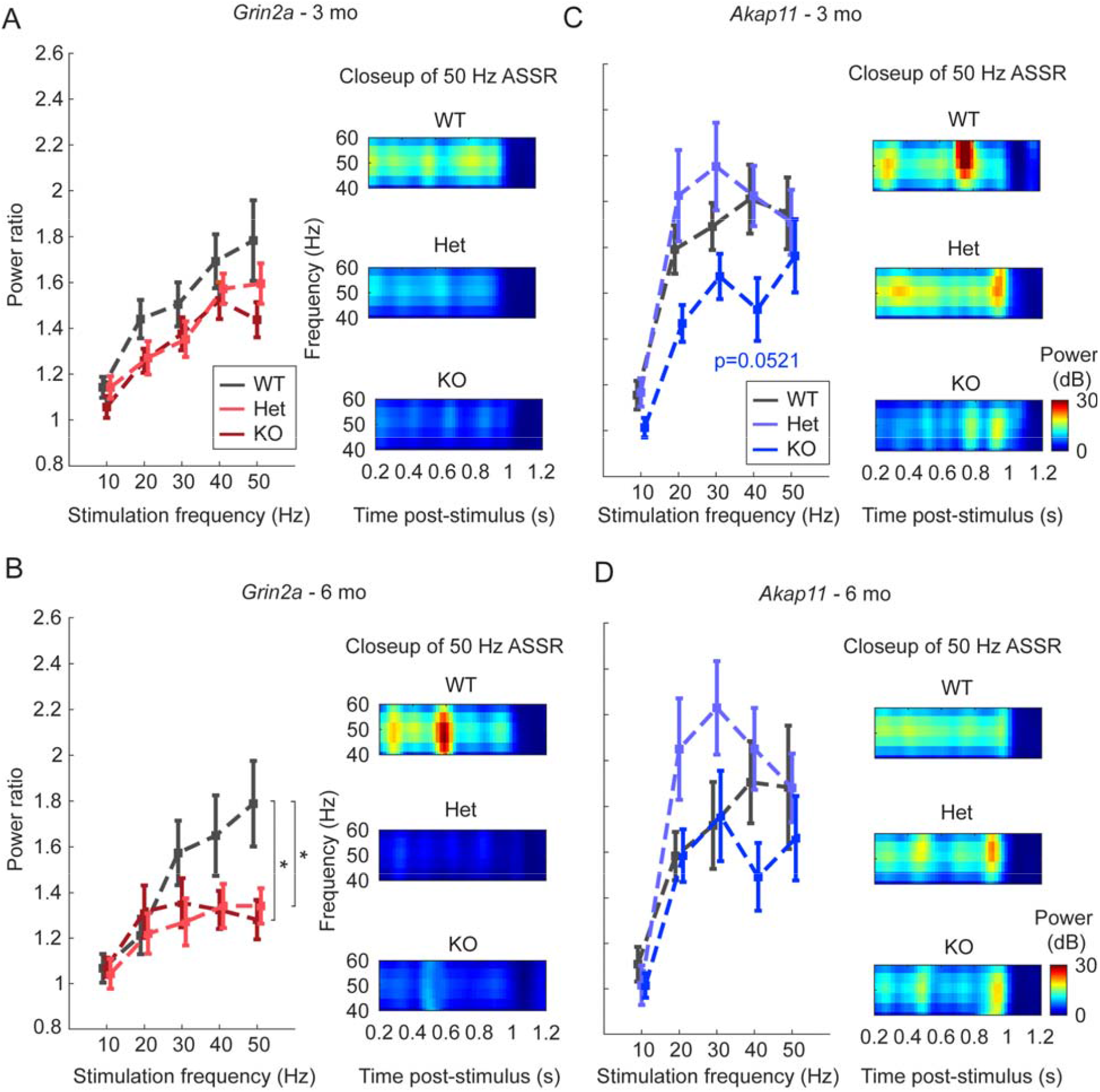
Auditory steady-state responses (ASSR) in *Grin2a* and *Akap11* mutants. **(A-D)** Left, power ratio quantification [Evoked/Baseline response] for 10-50 Hz auditory stimulation. Asterisks indicate significant differences in Het or KO vs. WT littermates (*p*<0.05). Error bars denote mean +/- standard error. Right, Z-scored power spectrogram visualization of the evoked response to 50 Hz stimulation, averaged across genotype. *n=*12 mice/group.

### *Grin2a*^*-/-*^ and *Akap11*^-/-^ mice show reduced responses to deviant auditory stimuli in the MMN paradigm

MMN (sometimes known as the “auditory oddball” paradigm) measures novelty-related responses to unexpected stimuli by comparing the event-related potentials in response to infrequent stimuli (deviant, 10% occurrence) vs. frequent stimuli (standard, 90% occurrence) that differ by at least one parameter (e.g. frequency, duration, or amplitude) [43, 44]. Schizophrenia patients display reduced responses to unexpected auditory stimuli, as reflected by the decreased amplitude of event-related potential components MMN and P3a, compared with healthy comparison subjects [24, 45]. These differential responses are thought to reflect deficits in sensory discrimination and novelty-related attention shifts, respectively. We found that *Grin2a* KO and *Akap11* KO exhibited altered responses to auditory stimuli during the MMN paradigm. *Grin2a*^-/-^ animals at 6 months (but not 3 months) exhibited reduced P1 (*p*=0.031) and reduced P3a (*p*=0.0099) peak amplitudes (∼30% and 40% decrease from WT levels, respectively) in response to deviant tones, while the N1 peak was increased in response to standard tones in these animals (∼35% increase from WT level, *p*=0.0217) (Figure 6A-B; Figure S6A-C). P1 amplitude in the difference waveform of 6-month-old *Grin2a* KO mice was reduced compared to WT and Het (Figure 6B), though this result was not statistically significant (*p*=0.1973) due to large variation in the data. *Akap11*^*-/-*^ animals showed a strongly reduced P3a amplitude in the difference waveform (response to deviant tones minus the response to standard tones) at 3 months (∼70% reduction vs. WT; *p*=0.0192), while *Akap11* heterozygous mutants had similar responses as their WT littermates (Figure 6C-D; S5D).

**Figure 6.**
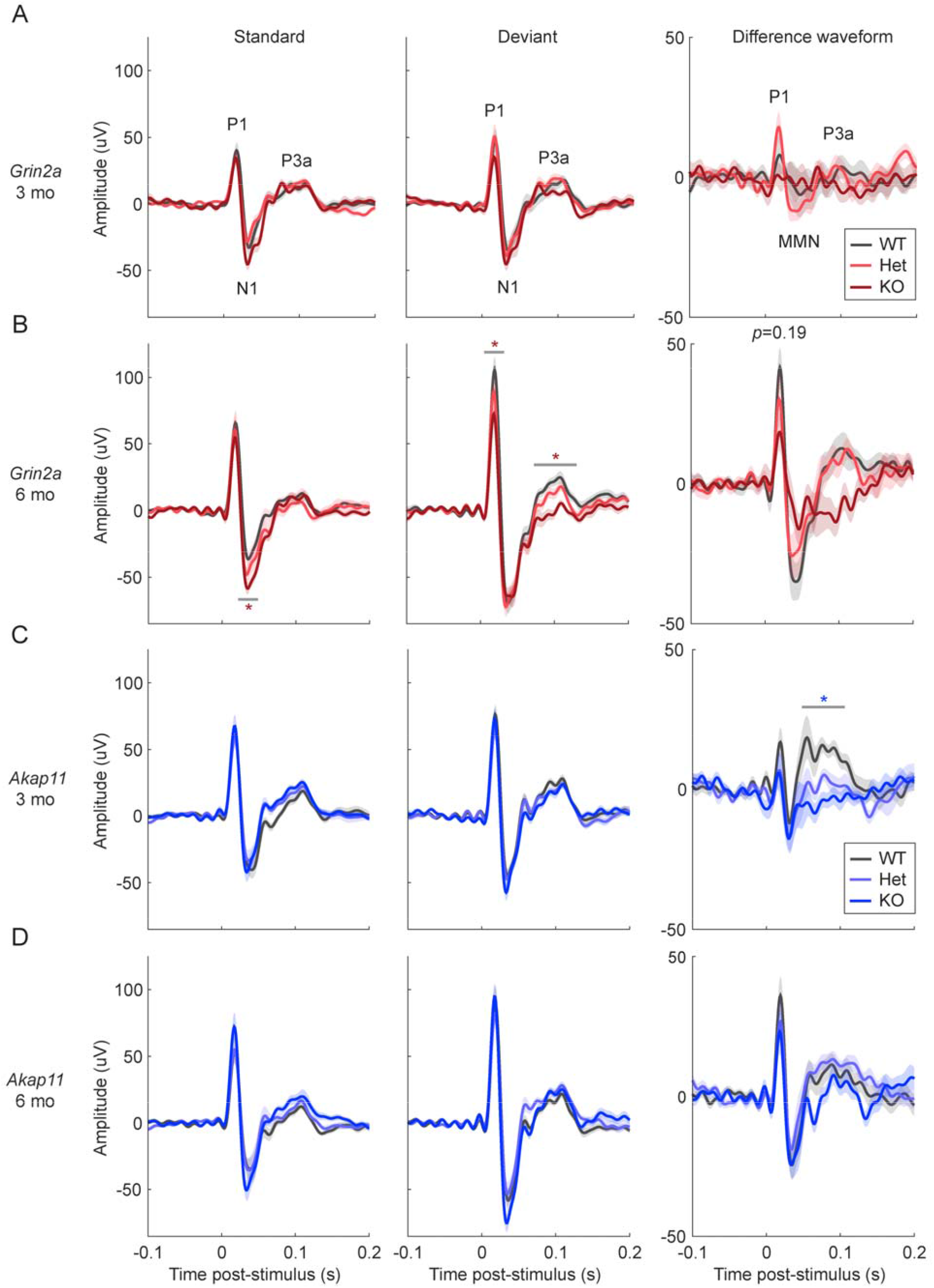
Mismatch negativity (MMN) responses in *Grin2a* and *Akap11* mutants. **(A-D)** Mean response to standard (*n=*900 trials), deviant (*n*=100 trials) and difference waveforms [response to deviant tones minus the response to standard tones], averaged across each genotype. Shading indicates mean +/- standard error. Red or blue asterisks indicate statistical differences in peak amplitude between WT and KO animals, which was assessed for the P1, N1/MMN and P3a peaks (as indicated on the first graphs). *p<*0.05, \*\**p<*0.01, \*\*\**p*<0.001; *n=*11-12 mice/group.

## DISCUSSION

Recent large-scale exome-sequencing efforts have uncovered rare genetic variants that confer high risk for schizophrenia, including *GRIN2A* and *AKAP11* [1, 2]. These relatively penetrant, loss-of-function mutations lend themselves to disease modeling in genetically modified mice. Besides their genetic validity, do *Grin2a* and *Akap11* mutant mice have any neurobiological phenotype that resembles the human disease (schizophrenia and/or bipolar disorder)? Using a neurophysiological assay (EEG) that is largely translatable between rodents and humans, we report that *Grin2a* and *Akap11* mutants show several key EEG abnormalities that are also observed in patients with schizophrenia and bipolar disorder, including increased resting gamma oscillations (Figure 3), attenuated 40-50 Hz ASSR (Figure 5), and abnormal MMN (in particular, altered P3a event-related potentials in the MMN paradigm) (Figure 6). These shared EEG abnormalities suggest that *Grin2a* and *Akap11* deficiencies impact, at least in part, the same brain networks and circuits that go awry in schizophrenia patients.

A major finding of our study is that *Grin2a* and *Akap11* mutants robustly show elevated resting gamma power (Figure 3), consistent with schizophrenia [25, 36, 37, 46] and to a lesser extent, bipolar disorder patients [47-49]. Gamma oscillations (defined here as 30-50 Hz) are generated by local cortical networks of fast-spiking, parvalbumin-positive (PV^+^) interneurons and excitatory pyramidal cells [41, 50, 51]. Deficiencies in PV^+^ interneurons have been implicated in postmortem studies of schizophrenia patients [52, 53], and resting gamma power is elevated in NMDAR hypofunction rodent models [54, 55], including mice lacking NMDAR transmission specifically in PV^+^ interneurons [51]. These and our findings are in line with the long-standing “glutamatergic hypothesis” that NMDAR dysfunction may be an underlying cause of schizophrenia [15, 56]. Indeed, NMDA receptors, including GRIN2A, are important for the function of PV^+^ interneurons [57], and undergo changes during development that may make these cells particularly sensitive to perturbation in psychiatric disease [58]. Importantly, even in the heterozygous state (which is the most relevant to human schizophrenia [1]), *Grin2a* mutations had a significant effect on resting gamma power (Figure 3B) and 40-50 Hz ASSR (Figure 5B) at 6 months. Overall, our findings suggest that *Grin2a* and *Akap11* deficiency affect convergent neural pathways, potentially involving circuits of PV^+^ interneurons and excitatory neurons, which are the cell types that drive gamma oscillations [59].

In contrast to gamma oscillations, which coordinate brain activity within local cortical networks, slow oscillations (0.5-1 Hz) preferentially establish long-range synchronization and are prominent during NREM sleep [60, 61]. Slow oscillations are altered in schizophrenia patients, although the direction of the effect is variable depending on brain region [62, 63]. The precise temporal organization of cortical slow oscillations, corticothalamic sleep spindles, and hippocampal sharp-wave ripples is thought to be important for sleep-dependent memory consolidation, which may be impaired in patients with schizophrenia [39, 64]. In our study, *Grin2a*^-/-^ mice exhibited increases in spontaneous power across multiple frequency bands, including slow oscillations (Figure 3), reminiscent of an “over-excitation” phenotype [65]. In contrast, broadband power increases (other than gamma) were not observed in *Akap11* mutants, and slow oscillations were reduced. Intriguingly, mutations in *GRIN2A*, but not *AKAP11*, are associated with certain epilepsy disorders [1, 3], and a drug that acts as a positive allosteric modulator of GRIN2A-containing NMDA receptors reduces spontaneous power (including slow oscillation power) in mouse models of Dravet syndrome and Alzheimer’s disease [32, 66]. These and our findings further substantiate a link between schizophrenia and epilepsy, which is affected by *GRIN2A*, but not *AKAP11* mutation.

The MMN test is a clinically translatable assay of sensory processing of familiar versus unexpected stimuli [43, 67]. The auditory event-related potential peak components P1, N1/MMN and P3a are thought to represent early detection, automatic sensory discrimination, and novelty-related attention shifts, respectively [68]. On average, schizophrenia [24, 69, 70] and bipolar disorder [71-73] patients tend to exhibit reduced MMN and P3a peak amplitudes, typically measured in the difference waveform (the responses to deviant stimuli minus the responses to standard stimuli). Interestingly, *Grin2a*^*-/-*^ (6 months) and *Akap11*^*-/-*^ mice (3 months) exhibited reduced P3a responses to deviant stimuli (Figure 6). The MMN peak in the difference waveform is reported to be reduced on average in schizophrenia and bipolar disorder patients; however, we found no significant changes in *Grin2a* or *Akap11* mutant mice, perhaps because of high variability in this metric [70, 71]. 6-month-old *Grin2a*^*-/-*^ mice exhibited increased N1 amplitude in response to standard tones (Figure 6B), potentially indicating lack of habituation to the repeated stimulus.

Sleep spindles (12-15 Hz) are another EEG feature typical of NREM sleep, generated by neurons across the cortex, thalamus, and thalamic reticular nucleus (TRN) [26]. Although rather heterogenous, patients with schizophrenia or bipolar disorder have on average a lower sleep spindle density relative to healthy comparison subjects [27, 35]. Despite sharing several other key features, the EEG phenotypes of *Grin2a* and *Akap11* mutants showed *opposite* and striking changes in sleep spindles. *Grin2a* mutants had elevated sleep spindle density, whereas *Akap11* mutants showed fewer spindles (Figure 4). In this context, it is noteworthy that both *Grin2a* and *Akap11* heterozygous mutants (which best model the human disease) exhibited a sleep spindle phenotype intermediate between homozygous knockouts and wild-type littermates. Because *Grin2a* and *Akap11* mutations have such disparate effects on spindle density, it is tempting to speculate that the heterogeneity of spindle density in human patients might arise in part from underlying genetic heterogeneity. The opposite effects of *Grin2a* and *Akap11* on sleep spindles implies that these genes differentially regulate the thalamocortical circuits responsible for spindle generation [39]. Although the precise neurophysiologic mechanisms remain to be discovered, our findings suggest that sleep spindles could serve as a biomarker separating different circuit mechanisms affected by *Grin2a* and *Akap11* loss-of-function, and potentially useful in stratifying the human disease population.

The convergent and divergent EEG features identified in this study highlight systems-level changes which may provide insight into the neurobiological pathways affected by loss of *Grin2a* and *Akap11*. Overall, the considerable overlap in EEG phenotypes between mouse mutants and human patients gives further credence to *Akap11*- and *Grin2a*-deficient mice as animal models of schizophrenia, particularly so as heterozygous mutants—SCHEMA mutations are found as heterozygotes in humans—tend to show abnormality intermediate between knockout and WT (e.g., *Grin2a*^+/-^ in gamma oscillation; *Grin2a*^+/-^ and *Akap11*^+/-^ in spindle density). Further investigation of the mechanistic foundations for these EEG changes, including transcriptomic and proteomic characterizations of *Grin2a* and *Akap11* mutants, will help elucidate which brain regions, cell types and molecular pathways contribute to the network-level dysfunctions of schizophrenia and bipolar disorder.

## ACKNOWLEDGEMENTS AND DISCLOSURES

We thank Nate Shepard for assisting with the mouse colony and Patrick Ihejirika for helping with data analysis. Research reported in this manuscript was supported by the Stanley Center for Psychiatric Research. Z.F. is supported by an Otto Hahn Fellowship of the Max Planck Society. M.S. serves on the Scientific Advisory Board of Biogen, Cerevel, Neumora, Vanqua Bio, and ArcLight Therapeutics. Results from this manuscript have been posted on the preprint server *bioRxiv*.

## REFERENCES

1. Singh T, Poterba T, Curtis D, Akil H, Al Eissa M, Barchas JD, et al. (2022): Exome sequencing identifies rare coding variants in 10 genes which confer substantial risk for schizophrenia. Nature: 10.1038/s41586-022-04556-w

2. Palmer DS, Howrigan DP, Chapman SB, Adolfsson R, Bass N, Blackwood D, et al. (2021): Exome sequencing in bipolar disorder reveals shared risk gene AKAP11 with schizophrenia. medRxiv: 2021.03.09.21252930.

3. Carvill GL, Regan BM, Yendle SC, O’Roak BR, Lozovaya N, Bruneau N, et al. (2013): GRIN2A mutations cause epilepsy-aphasia spectrum disorders. Nature Genetics 45(9): 1073–6.

4. Salmi M, Bolbos R, Bauer S, Minlebaev M, Burnashev N, Szepetowski P (2018): Transient microstructural brain anomalies and epileptiform discharges in mice defective for epilepsy and language-related NMDA receptor subunit gene Grin2a. Epilepsia 59(10): 1919–1930.

5. The Schizophrenia Working Group of the Psychiatric Genomics Consortium, Ripke S, Neale BM, Corvin A, Walters JTR, Farh K, et al. (2014): Biological insights from 108 schizophrenia-associated genetic loci. Nature 511(7510): 421–7.

6. The Schizophrenia Working Group of the Psychiatric Genomics Consortium, Ripke S, Walters JTR, O’Donovan MC (2022): Mapping genomic loci prioritises genes and implicates synaptic biology in schizophrenia. Nature: 10.1038/s41586-022-04434-5.

7. Dennison CA, Legge SE, Pardiñas AF, Walters JTR (2020): Genome-wide association studies in schizophrenia: Recent advances, challenges and future perspective. Schizophrenia Research 217: 4–12.

8. Marshall CR, Howrigan DP, Merico D, Thiruvahindrapuram B, Wu W, Greer DS, et al. (2017): Contribution of copy number variants to schizophrenia from a genome-wide study of 41,321 subjects. Nature Genetics 49(1): 27–35.

9. Genovese G, Fromer M, Stahl EA, Ruderfer DM, Chambert K, Landén M, et al. (2016): Increased burden of ultra-rare protein-altering variants among 4,877 individuals with schizophrenia. Nature Neuroscience 19(11): 1433–1441.

10. Singh T, Walters JTR, Johnstone M, Curtis D, Suvisaari J, Torniainen M, et al. (2017): The contribution of rare variants to risk of schizophrenia in individuals with and without intellectual disability. Nature Genetics 49(8): 1167–1173.

11. Rivas MA, Pirinen M, Conrad DF, Lek M, Tsang EK, Karczewski KJ, et al. (2015): Human genomics. Effect of predicted protein-truncating genetic variants on the human transcriptome. Science 348(6235): 666–9.

12. Miyamoto S, Miyake N, Jarskog LF, Fleischhacker WW, Lieberman JA (2012): Pharmacological treatment of schizophrenia: a critical review of the pharmacology and clinical effects of current and future therapeutic agents. Molecular Psychiatry 17(12): 1206–27.

13. Paoletti P, Bellone C, Zhou Q (2013): NMDA receptor subunit diversity: impact on receptor properties, synaptic plasticity and disease. Nature Reviews Neuroscience 14(6): 383–400.

14. Zhou Q, Sheng M (2013): NMDA receptors in nervous system diseases. Neuropharmacology 74: 69–75.

15. Moghaddam B, Javitt D (2012): From revolution to evolution: the glutamate hypothesis of schizophrenia and its implication for treatment. Neuropsychopharmacology, 37(1): 4–15.

16. Solmi M, Radua J, Olivola M, Croce E, Soardo L, Salazar de Pablo G, et al. (2012): Age at onset of mental disorders worldwide: large-scale meta-analysis of 192 epidemiological studies. Molecular Psychiatry 27: 281–295.

17. Sheng M, Cummings J, Roldan LA, Jan YN, Jan LY (1994): Changing subunit composition of heteromeric NMDA receptors during development of rat cortex. Nature 368(6467): 144–7.

18. Ohi K, Shimada T, Nitta Y, Kihara H, Okubo H, Uehara T, et al. (2016): Specific gene expression patterns of 108 schizophrenia-associated loci in cortex. Schizophrenia Research 174(1-3): 35-38.

19. Deng Z, Li X, Ramirez MB, Purtell K, Choi I, Lu J, et al. (2021): Selective autophagy of AKAP11 activates cAMP/PKA to fuel mitochondrial metabolism and tumor cell growth. PNAS 118(14): e2020215118.

20. Varjosalo M, Keskitalo S, Van Drogen A, Nurkkala H, Vichalkovski A, Aebersold R, et al. (2013): The protein interaction landscape of the human CMGC kinase group. Cell Reports 3(4): 1306–20.

21. Freland L, Beaulieu JM (2012): Inhibition of GSK3 by lithium, from single molecules to signaling networks. Frontiers in Molecular Neuroscience 5: p. 14.

22. Uhlhaas PJ, Singer W (2010): Abnormal neural oscillations and synchrony in schizophrenia. Nature Reviews Neuroscience 11(2): 100–13.

23. Gonzalez-Burgos G, Lewis DA (2012): NMDA receptor hypofunction, parvalbumin-positive neurons, and cortical gamma oscillations in schizophrenia. Schizophrenia Bulletin 38(5): 950–7.

24. Light GA, Swerdlow NR, Thomas ML, Calkins ME, Green MF, Greenwood TA, et al. (2015): Validation of mismatch negativity and P3a for use in multi-site studies of schizophrenia: characterization of demographic, clinical, cognitive, and functional correlates in COGS-2. Schizophrenia Research 163(1-3): 63–72.

25. Thuné H, Recasens M, Uhlhaas PJ (2016): The 40-Hz Auditory Steady-State Response in Patients With Schizophrenia: A Meta-analysis. JAMA Psychiatry 73(11): 1145–1153.

26. Manoach DS, Pan JQ, Purcell SM, Stickgold R (2016): Reduced Sleep Spindles in Schizophrenia: A Treatable Endophenotype That Links Risk Genes to Impaired Cognition? Biological Psychiatry 80(8): 599–608.

27. Kozhemiako N, Wang J, Jiang C, Wang L, Gai G, Zou K, et al. (2021): Non-rapid eye movement sleep and wake neurophysiology in schizophrenia. bioRxiv: 2021.12.13.472475.

28. Kadotani H, Hirano T, Masugi M, Nakamura K, Nakao K, Katsuki M, et al. (1996): Motor discoordination results from combined gene disruption of the NMDA receptor NR2A and NR2C subunits, but not from single disruption of the NR2A or NR2C subunit. The Journal of Neuroscience 16(24): 7859–67.

29. Whiting JL, Ogier L, Forbush KA, Bucko P, Gopalan J, Seternes O, et al. (2016): AKAP220 manages apical actin networks that coordinate aquaporin-2 location and renal water reabsorption. PNAS 113(30): E4328–37.

30. Walsh RN, Cummins RA (1976): The Open-Field Test: a critical review. Psychological Bulletin 83(3): 482–504.

31. Seibenhener ML, Wooten MC (2015): Use of the Open Field Maze to measure locomotor and anxiety-like behavior in mice. Journal of Visualized Experiments 96: e52434.

32. Hanson JE, Ma K, Elstrott J, Weber M, Saillet S, Khan AS, et al. (2020): GluN2A NMDA Receptor Enhancement Improves Brain Oscillations, Synchrony, and Cognitive Functions in Dravet Syndrome and Alzheimer’s Disease Models. Cell Reports 30(2): 381–396.

33. Wulff K, Dijk D, Middleton B, Foster RG, Joyce EM (2012): Sleep and circadian rhythm disruption in schizophrenia. British Journal of Psychiatry 200(4): 308–16.

34. Chan M, Chung K, Yung K, Yeung W (2017): Sleep in schizophrenia: A systematic review and meta-analysis of polysomnographic findings in case-control studies. Sleep Medicine Reviews 32: 69–84.

35. Zangani C, Casetta C, Saunders AS, Donati F, Maggioni E, D’Agostino A (2020): Sleep abnormalities across different clinical stages of Bipolar Disorder: A review of EEG studies. Neuroscience & Biobehavioral Reviews 118: 247–257.

36. Tekell JL, Hoffmann R, Hendrickse W, Greene RW, Rush AJ, Armitage R (2005): High frequency EEG activity during sleep: characteristics in schizophrenia and depression. Clinical EEG and Neuroscience 36(1): 25–35.

37. Tanaka-Koshiyama K, Koshiyama D, Miyakoshi M, Joshi YB, Molina JL, Sprock J, et al. (2020): Abnormal Spontaneous Gamma Power Is Associated With Verbal Learning and Memory Dysfunction in Schizophrenia. Frontiers in Psychiatry 11: 832.

38. Ferrarelli F (2015): Sleep in patients with schizophrenia. Current Sleep Medicine Reports 1(2): 150–156.

39. Manoach DS, Stickgold R (2019): Abnormal Sleep Spindles, Memory Consolidation, and Schizophrenia. Annual Review of Clinical Psychology 15: 451–479.

40. O’Donnell BF, Vohs JL, Krishnan GP, Rass O, Hetrick WP, Morzorati SL (2013): The auditory steady-state response (ASSR): a translational biomarker for schizophrenia. Supplements to Clinical Neurophysiology 62: 101–12.

41. Sun C, Zhou P, Wang C, Fan Y, Tian Q, Dong F, et al. (2018): Defects of Gamma Oscillations in Auditory Steady-State Evoked Potential of Schizophrenia. Shanghai Archives of Psychiatry 30(1): 27–38.

42. Sugiyama S, Ohi Z, Kuramitsu A, Takai K, Muto Y, Taniguchi T, et al. (2021): The Auditory Steady-State Response: Electrophysiological Index for Sensory Processing Dysfunction in Psychiatric Disorders. Frontiers in Psychiatry 12: 644541.

43. Tada M, Kirihara K, Mizutani S, Uka T, Kunii N, Koshiyama D, et al. (2019): Mismatch negativity (MMN) as a tool for translational investigations into early psychosis: A review. International Journal of Psychophysiology 145: p. 5–14.

44. Herzog L, Salehi KL, Bohon KS, Wiest MC (2014): Prestimulus frontal-parietal coherence predicts auditory detection performance in rats. Journal of Neurophysiology 111(10): 1986–2000.

45. Kaur M, Battisti RA, Ward PB, Ahmed A, Hickie IB, Hermens DF (2011): MMN/P3a deficits in first episode psychosis: comparing schizophrenia-spectrum and affective-spectrum subgroups. Schizophrenia Research 130(1-3): 203-9.

46. Sivarao DV, Chen P, Senapati A, Yang Y, Fernandes A, Benitex Y (2016): 40 Hz Auditory Steady-State Response Is a Pharmacodynamic Biomarker for Cortical NMDA Receptors. Neuropsychopharmacology 41(9): 2232–40.

47. Narayanan B, O’Neil K, Berwise C, Stevens MC, Calhoun VD, Clementz BA (2014): Resting state electroencephalogram oscillatory abnormalities in schizophrenia and psychotic bipolar patients and their relatives from the bipolar and schizophrenia network on intermediate phenotypes study. Biological Psychiatry 76(6): 456–65.

48. Venables NC, Bernat EM, Sponheim SR (2009): Genetic and disorder-specific aspects of resting state EEG abnormalities in schizophrenia. Schizophrenia Bulletin 35(4): 826–39.

49. Arikan MK, Günver G, İlhan R, Öksüz O, Metin B (2021): Gamma oscillations predict treatment response to aripiprazole in bipolar disorder. Journal of Affective Disorders 294: 159–162.

50. Buzsaki G, Wang XJ (2012): Mechanisms of gamma oscillations. Annual Review of Neuroscience 35: 203–25.

51. Carlén M, Meletis K, Siegle JH, Cardin JA, Futai K, Vierling-Claassen D (2011): A critical role for NMDA receptors in parvalbumin interneurons for gamma rhythm induction and behavior. Molecular Psychiatry 17(5): 537–48.

52. Nakazawa K, Zsiros V, Jiang Z, Nakao K, Kolata S, Zhang S, et al. (2012): GABAergic interneuron origin of schizophrenia pathophysiology. Neuropharmacology 62(3): 1574–83.

53. Lewis DA, Curley AA, Glausier JR, Volk DW (2012): Cortical parvalbumin interneurons and cognitive dysfunction in schizophrenia. Trends in Neuroscience 35(1): 57–67.

54. Nakao K, Singh M, Sapkota K, Hagler BC, Hunter RN, Raman C, et al. (2020): GSK3beta inhibition restores cortical gamma oscillation and cognitive behavior in a mouse model of NMDA receptor hypofunction relevant to schizophrenia. Neuropsychopharmacology 45(13): 2207–2218.

55. Gandal MJ, Sisti J, Klook K, Ortinski PI, Leitman V, Liang Y, et al. (2012): GABAB-mediated rescue of altered excitatory-inhibitory balance, gamma synchrony and behavioral deficits following constitutive NMDAR-hypofunction. Translational Psychiatry, 2: e142.

56. Uno Y, Coyle JT (2019): Glutamate hypothesis in schizophrenia. Psychiatry and Clinical Neurosciences 73(5): 204–215.

57. Kinney JW, Davis CN, Tabarean I, Conti B, Bartfai T, Behrens MM (2006): A specific role for NR2A-containing NMDA receptors in the maintenance of parvalbumin and GAD67 immunoreactivity in cultured interneurons. The Journal of Neuroscience 26(5): 1604–15.

58. Wang HX, Gao WJ (2009): Cell type-specific development of NMDA receptors in the interneurons of rat prefrontal cortex. Neuropsychopharmacology 34(8): 2028–40.

59. Cardin JA, Carlén M, Meletis K, Knoblich U, Zhang F, Deisseroth K, et al. (2009): Driving fast-spiking cells induces gamma rhythm and controls sensory responses. Nature 459(7247): 663–7.

60. D’Agostino A, Castelnovo A, Cavallotti S, Casetta C, Marcatili M, Gambini O, et al. (2018): Sleep endophenotypes of schizophrenia: slow waves and sleep spindles in unaffected first-degree relatives. Schizophrenia, 4(1): 2.

61. Moran LV, Hong LE (2011): High vs low frequency neural oscillations in schizophrenia. Schizophrenia Bulletin 37(4): 659–63.

62. Zhang Y, Quinones GM, Ferrarelli F (2020): Sleep spindle and slow wave abnormalities in schizophrenia and other psychotic disorders: Recent findings and future directions. Schizophrenia Research, 221: 29–36.

63. Ritter PS, Schwabedal J, Brandt M, Schrempf W, Brezan F, Krupka A, et al. (2018): Sleep spindles in bipolar disorder - a comparison to healthy control subjects. Acta Psychiatrica Scandinavica 138(2): 163–172.

64. Herzog LE, Katz DB, Jadhav SP (2020): Refinement and Reactivation of a Taste-Responsive Hippocampal Network. Current Biology 30(7): 1306–1311.

65. Wang W (2021): The broadband power shifts in entorhinal EEG are related to the firing of grid cells. Heliyon, 7(1): e06087.

66. Hackos DH, Hanson JE (2017): Diverse modes of NMDA receptor positive allosteric modulation: Mechanisms and consequences. Neuropharmacology 112(Pt A): 34–45.

67. Todd J, Harms L, Schall U, Michie PT (2013): Mismatch negativity: translating the potential. Frontiers in Psychiatry 4: 171.

68. Rissling AJ, Light GA (2010): Neurophysiological measures of sensory registration, stimulus discrimination, and selection in schizophrenia patients. Current Topics in Behavioral Neuroscience 4: 283–309.

69. Nagai T, Kirihara K, Tada M, Koshiyama D, Koike S, Suga M, et al. (2017): Reduced Mismatch Negativity is Associated with Increased Plasma Level of Glutamate in First-episode Psychosis. Scientific Reports 7(1): 2258.

70. Jahshan C, Wynn J, Mathis KI, Altshuler LL, Glahn DC, Green MF (2012): Cross-diagnostic comparison of duration mismatch negativity and P3a in bipolar disorder and schizophrenia. Bipolar Disorders 14(3): 239–48.

71. Umbricht D, Vyssotki D, Latanov A, Nitsch R, Lipp H (2005): Deviance-related electrophysiological activity in mice: is there mismatch negativity in mice? Clinical Neurophysiology 116(2): 353–63.

