## Supplemental Information for "Mouse mutants in schizophrenia risk genes *GRIN2A* and *AKAP11* show EEG abnormalities in common with schizophrenia patients"

### SUPPLEMENTAL MATERIALS AND METHODS

#### EEG recording

Sleep/wake: Signals were recorded for 24 hours from the onset of the dark phase, ZT12 (6 pm EDT or 7 pm EST).

Auditory steady-state responses (ASSR): MATLAB was used to generate 1 s auditory click trains of 10-50 Hz through the audio speaker (GTO629, JBL by Harman) at ~70 dB. A CompactDAQ system (cDAQ-9171, National Instruments) was used to generate a TTL pulse with each click train sent to Sirenia software to determine stimulus timing throughout the test. 20 randomized trials of each stimulus were delivered for a total of 100 trials lasting 25 minutes.

Mismatch negativity (MMN): MATLAB was used to generate 100 ms tones of either 6000 or 8000 Hz through the audio speaker (GTO629, JBL by Harman) at ~70 dB. A CompactDAQ system (cDAQ-9171, National Instruments) was used to generate a TTL pulse with each click train that was sent to Sirenia software to determine stimulus timing throughout the test. Each test consisted of 1000 randomized trials of 900 standard (6000 Hz) stimuli and 100 deviant (8000 Hz) stimuli. To control for animals' differential response to pitch, a "flip-flop control" test (using 8000 Hz standard and 6000 Hz deviant tones) was conducted on a different day, and the response to the same tone (8000 Hz) as deviant vs. standard was compared.

**EEG analysis**

Sleep/wake: Sleep state classification (NREM, REM or Wake) was performed on 10 s epochs of EEG/EMG data using a machine learning model developed by LW. The model is developed using the light gradient boosting machine (LightGBM), which is an ensemble model based on the decision tree, built using well-established EEG/EMG hallmarks of behavioral state (e.g., EMG amplitude, delta oscillation amplitude, delta/theta ratio). The LightGBM model achieves an overall accuracy of 95.2% and a Cohen's kappa of 0.91 (a measurement of the inter-rater reliability that accounts for the agreement rate by chance; kappa values greater than 0.8 are considered as nearly perfectly in agreement [1]), which is on par with the performance of human experts reading the EEG traces manually. Wake and NREM had higher f1-scores (the harmonic mean of precision and recall) than REM; wake = 0.96, NREM = 0.95, and REM = 0.85. LUNA software for EEG analysis (http://zzz.bwh.harvard.edu/luna) was used to compute stage duration, sleep onset (defined as % time asleep during the first hour of the light cycle), bout length and number, sleep spindle density, absolute and relative power for each band (slow: 0.5-1 Hz, delta: 1-4 Hz, theta: 4-8 Hz, alpha: 8-12 Hz, sigma: 12-15 Hz, beta: 15-30 Hz, gamma: 30-50 Hz) separately for the light and dark cycle. Measurements taken during NREM/REM sleep were analyzed during the light cycle, where mice are predominantly asleep. One 3-month-old WT Grin2a animal was excluded from NREM/REM power analyses due to excessive line noise during the light cycle. Measurements taken during wake were analyzed during the dark cycle, where mice are predominantly awake. Periods of quiet wake were defined as 10 s epochs where the mean EMG amplitude during wake did not exceed 3 standard deviations above or below the mean EMG amplitude during NREM sleep (the mouse is presumed to be immobile). Statistical differences between WT, Het and KO animals were computed using one-way ANOVAs with post hoc Tukey-Kramer tests for multiple comparisons.

Auditory steady-state responses (ASSR): ASSR were analyzed using custom MATLAB scripts to quantify the evoked response to different frequency auditory stimuli [2]. Trials were segmented into 2 s windows (0.5 s before stimulus onset to 1.5 s after stimulus onset) as detected from the TTL pulse. Successive trials of the same stimulus were removed from the analysis, as well as large-amplitude (>1800 uV) trials containing movement-related artifacts. EEG power was evaluated corresponding to the stimulus frequency (10-50 Hz) using the *pwelch* algorithm (0.5 s window, 0.25 s overlap) during the pre-stimulus (0.5 s before stimulus onset) and stimulus-evoked (0.2 to 1 s following stimulus onset) periods. The average power ratio (evoked/pre-stimulus power) for each stimulus type was statistically compared between genotypes using one-way ANOVAs with post hoc Tukey-Kramer tests for multiple comparisons. Z-scored average power spectrograms of the evoked responses (0.2 to 1 s following stimulus onset) for each group were also plotted for visualization using the spectrogram algorithm.

Mismatch negativity (MMN): The peak components of event-related potentials (ERPs) generated from the MMN test (P1, N1/MMN, and P3a) were analyzed using custom MATLAB scripts [3]. P1 refers to the first positive peak of the ERP. N1 and MMN both refer to the first negative peak of the ERP (deemed N1 in the response to standard or deviant tones, and MMN in the difference waveform). P3a refers to the second positive peak in the ERP. Trials were segmented into 1 s windows (0.4 s before stimulus onset to 0.6 s after stimulus onset) as detected from the TTL pulse. Trials were normalized to the baseline response (average voltage across the trial) and large-amplitude (>1800 uV) trials containing movement artifacts were removed from the analysis. The average evoked potentials from standard trials preceding the deviant, deviant trials, and the difference waveform (deviant-standard) were plotted for each genotype for visualization. The peak amplitude of the P1 (maximum peak detected 0-0.05 s post-stimulus), N1/MMN (minimum peak detected 0-0.1 s post-stimulus) and P3a (maximum peak detected 0.05-0.15 s post-stimulus) components were calculated for each animal for statistical comparison across groups using one-way ANOVAs with post hoc Tukey-Kramer tests for multiple comparisons.

### REFERENCES

1. McHugh ML (2012): Interrater reliability: the kappa statistic. *Biochemia Medica* 22(3): 276-82.

2. O'Donnell BF, Vohs JL, Krishnan GP, Hetrick WP, Morzorati SL (2013): The auditory steady-state response (ASSR): a translational biomarker for schizophrenia. *Supplements to Clinical Neurophysiology* 62: 101-12.

3. Tada M, Kirihara K, Mizutani SD, Uka T, Kunii N, Koshiyama D, *et al.* (2019): Mismatch negativity (MMN) as a tool for translational investigations into early psychosis: A review. *International Journal of Psychophysiology* 145: 5-14.

### SUPPLEMENTAL FIGURES

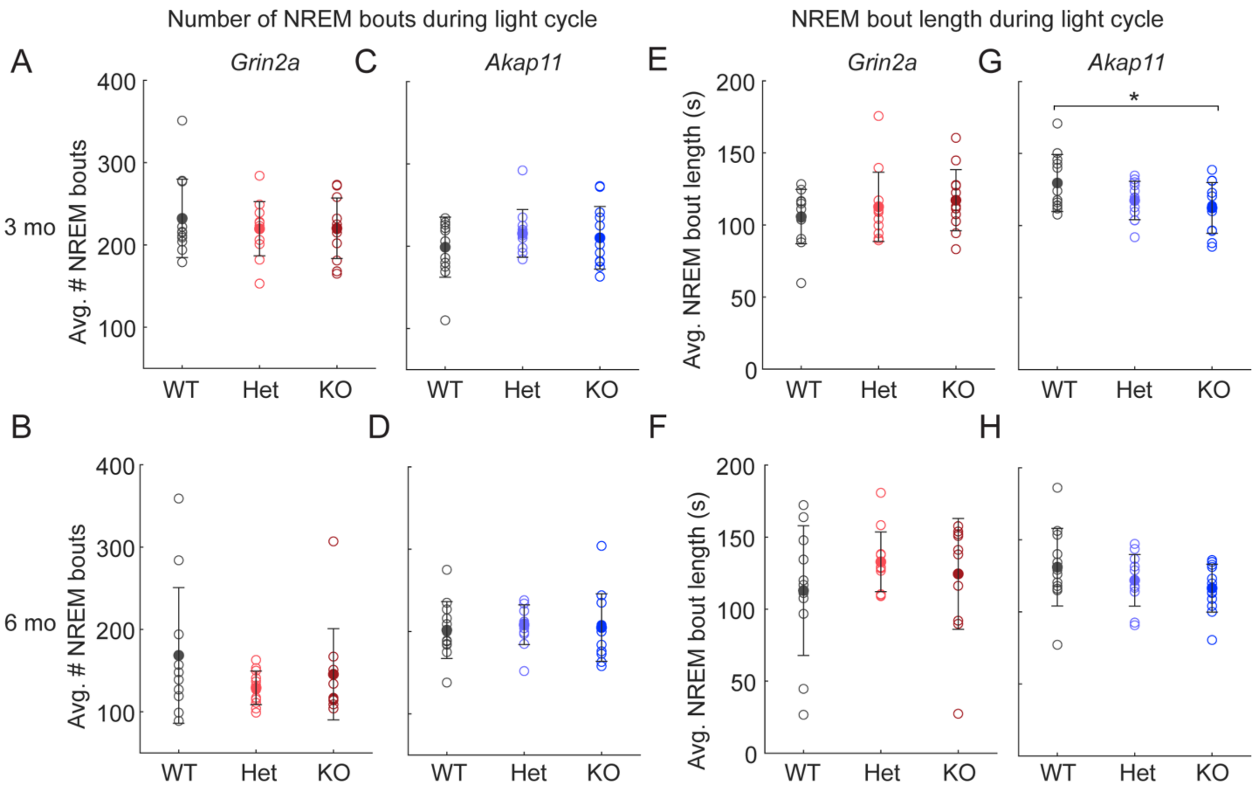


**Figure S1. Sleep fragmentation patterns of *Grin2a* and *Akap11* mutant mice.** (A-D) Average number of NREM bouts during the light cycle. *Grin2a* and *Akap11* mutants both exhibited similar bout numbers as their WT littermates, suggesting the NREM deficiencies found in *Akap11*$^{-/-}$ mice (see Figure 2) are not due to sleep fragmentation. (E-H) Average NREM bout length during the light cycle. At 3 months, *Akap11*$^{-/-}$ mice exhibited shorter NREM bout lengths than their WT littermates (p=0.0478). Error bars denote mean +/- standard error; p<0.05, **p<0.01, ***p<0.001; n=11-12 mice/group.

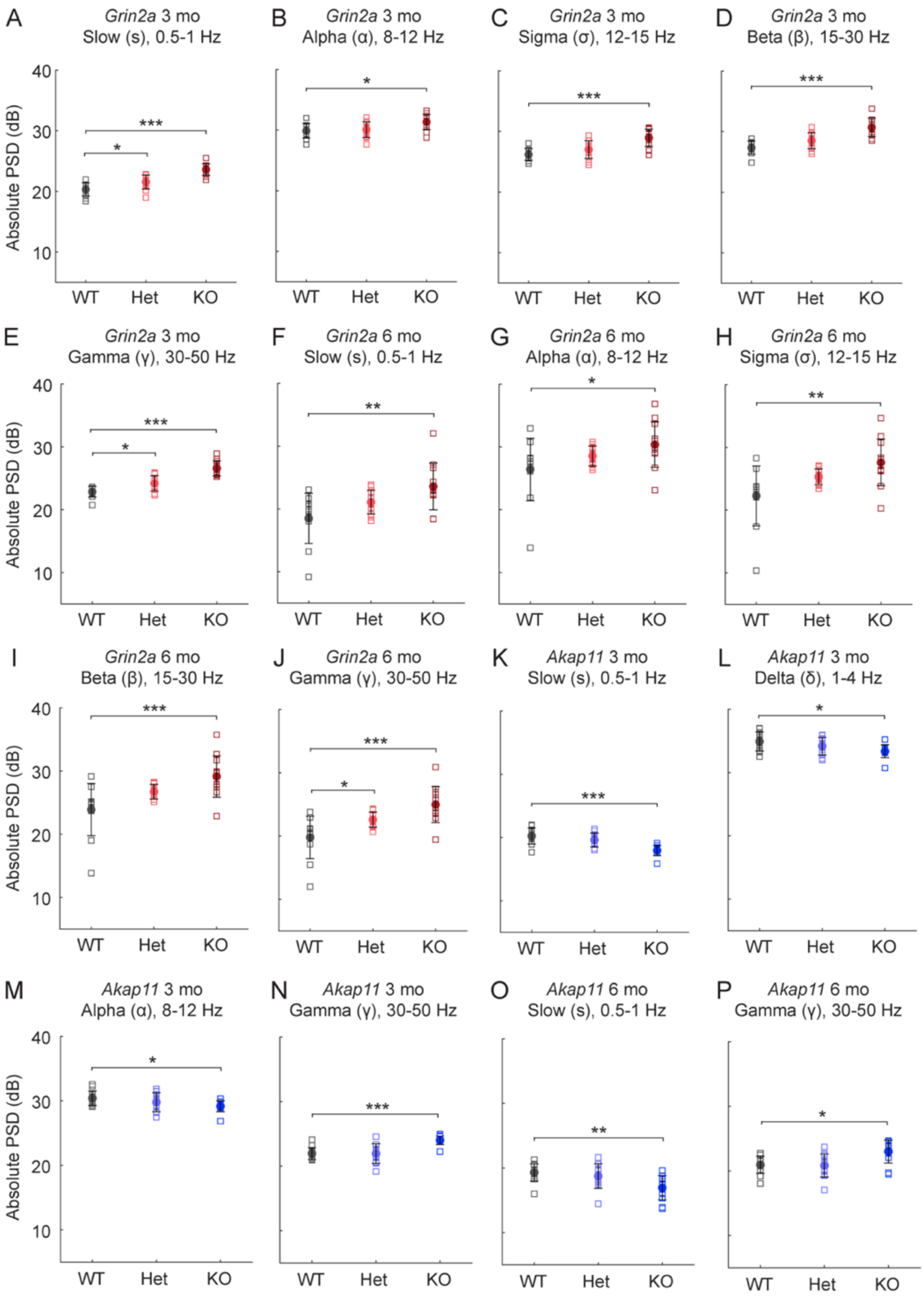

A
*Grin2a* 3 mo
Slow (s), 0.5-1 Hz
B
*Grin2a* 3 mo
Alpha (α), 8-12 Hz
C
*Grin2a* 3 mo
Sigma (σ), 12-15 Hz
D
*Grin2a* 3 mo
Beta (β), 15-30 Hz
E
*Grin2a* 3 mo
Gamma (γ), 30-50 Hz
F
*Grin2a* 6 mo
Slow (s), 0.5-1 Hz
G
*Grin2a* 6 mo
Alpha (α), 8-12 Hz
H
*Grin2a* 6 mo
Sigma (σ), 12-15 Hz
I
*Grin2a* 6 mo
Beta (β), 15-30 Hz
J
*Grin2a* 6 mo
Gamma (γ), 30-50 Hz
K
*Akap11* 3 mo
Slow (s), 0.5-1 Hz
L
*Akap11* 3 mo
Delta (δ), 1-4 Hz
M
*Akap11* 3 mo
Alpha (α), 8-12 Hz
N
*Akap11* 3 mo
Gamma (γ), 30-50 Hz
O
*Akap11* 6 mo
Slow (s), 0.5-1 Hz
P
*Akap11* 6 mo
Gamma (γ), 30-50 Hz
Absolute PSD (dB)
10
20
30
40
WT
Het
KO

**Figure S2. Absolute PSD of brain oscillations in *Grin2a* and *Akap11* mutant mice during NREM sleep.** Absolute power for 3- and 6-month *Grin2a* (A-J) and *Akap11* (K-P) mutants in NREM sleep during the light cycle. Absolute PSD was computed for each of the following frequency bands: slow (s), 0.5-1 Hz; delta (δ), 1-4 Hz; theta (θ), 4-8 Hz; alpha (α), 8-12 Hz; sigma (σ), 12-15 Hz; beta (β), 15-30 Hz; gamma (γ), 30-50 Hz. Quantification of significant differences (see Figure 3) are shown here. Error bars show mean +/- standard error; $p<0.05$, **$p<0.01$, ***$p<0.001$; n=11-12 mice/group.

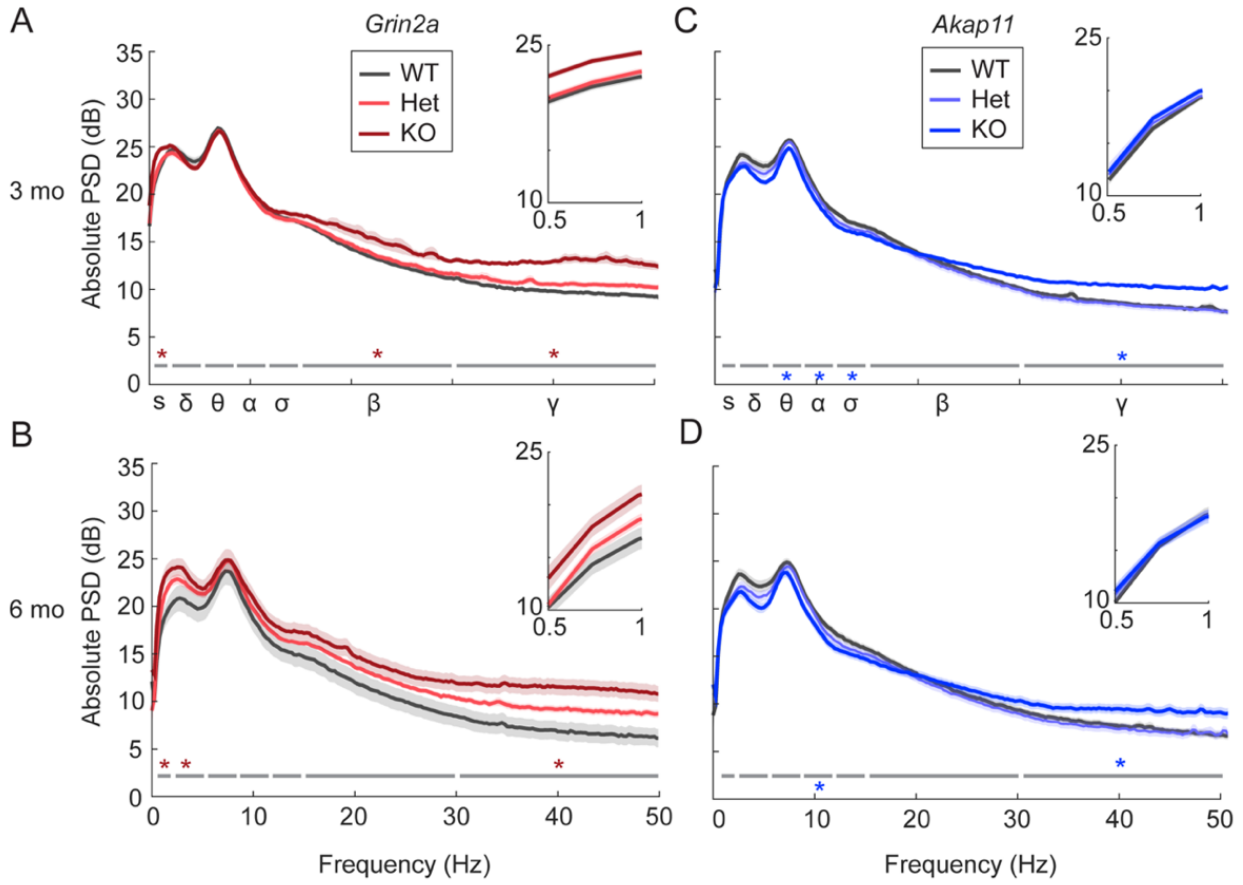


**Figure S3. Power spectral analysis of brain oscillations in *Grin2a* and *Akap11* mutant mice during REM sleep.** (A-D) Absolute REM power spectra for 3- and 6-month *Grin2a* and *Akap11* mutants during the light cycle. Light and dark red/blue stars indicate oscillations for which +/- and -/- mice, respectively, differed significantly from WT littermates (p<0.05) in each of the frequency bands: slow (s), 0.5-1 Hz; delta (δ), 1-4 Hz; theta (θ), 4-8 Hz; alpha (α), 8-12 Hz; sigma (σ), 12-15 Hz; beta (β), 15-30 Hz; gamma (γ), 30-50 Hz; with stars above or below the gray line indicating significant increases or decreases in power, respectively. *Grin2a*$^{-/-}$ (3 months: p=2.99e-07; 6 months: p=8.29e-04) and *Akap11*$^{-/-}$ (3 months: p=2.87e-04; 6 months: p=0.0435) mice exhibited increased gamma oscillations, similar to NREM sleep (Figure 3). *Grin2a*$^{-/-}$ animals additionally exhibited increases in slow (3 months: p=7.95e-07; 6 months: p=0.0056), beta (3 months: p=0.0195) and delta oscillations (6 months: p=0.0406), while *Akap11*$^{-/-}$ animals had reduced theta (3 months: p=0.0138), alpha (3 months: p=0.0036; 6

months: p=0.0473) and sigma (3 months: p=0.0397) oscillations. Shading indicates mean +/- standard error. Insets show magnified view of the graph in the slow oscillation range (0.5-1 Hz). n=11-12 mice/group.

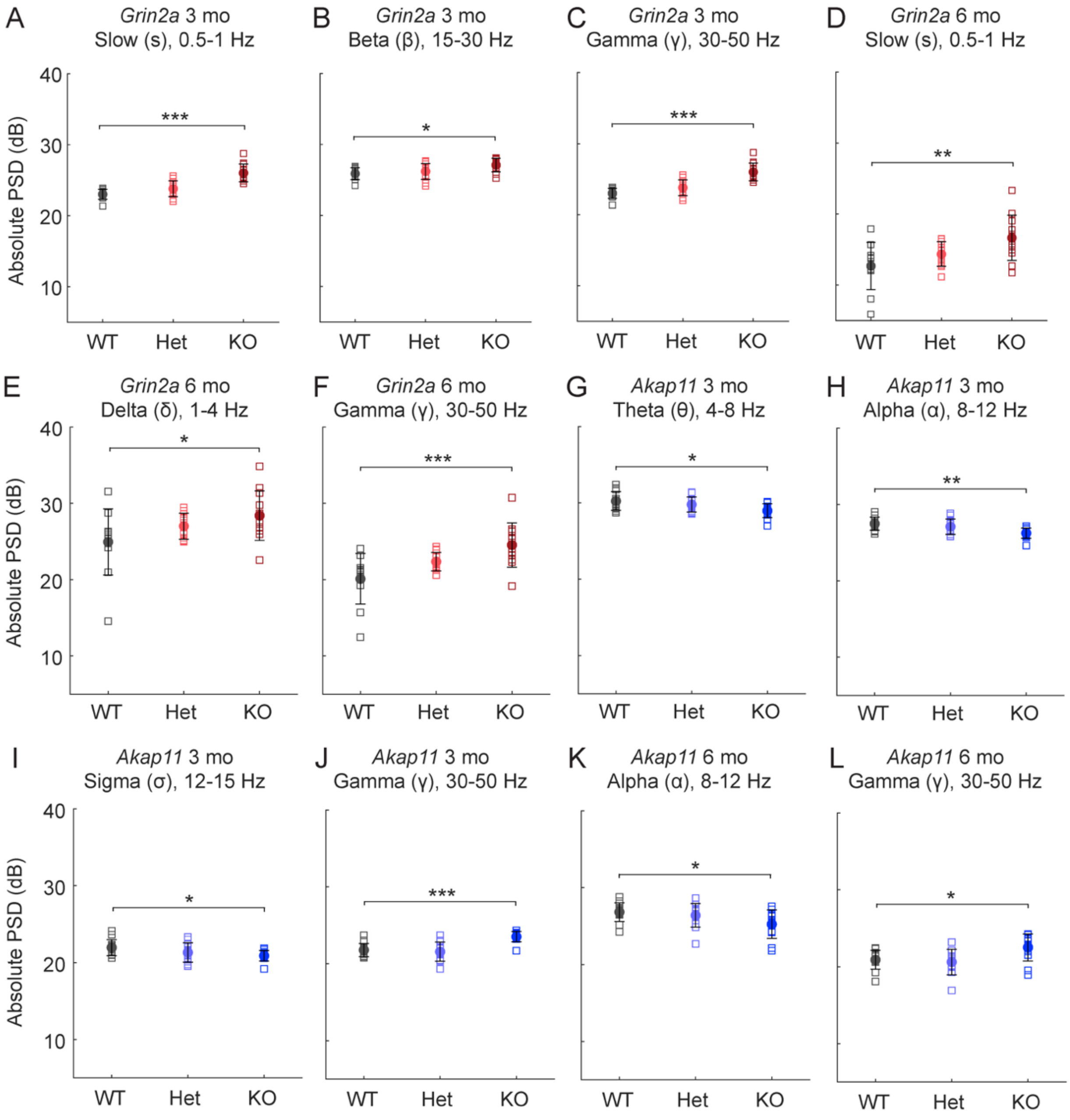


**Figure S4. Absolute PSD of brain oscillations in *Grin2a* and *Akap11* mutant mice during REM sleep**. Absolute power for 3- and 6-month *Grin2a* (A-J) and *Akap11* (K-P) mutants in REM sleep during the light cycle. Absolute PSD was computed for each of the following frequency bands: slow (s), 0.5-1 Hz; delta (δ), 1-4 Hz; theta (θ), 4-8 Hz; alpha (α), 8-12 Hz; sigma (σ), 12-15 Hz; beta (β), 15-30 Hz; gamma (γ), 30-50 Hz. Quantification of significant differences (see

Figure S3) are shown here. Error bars show mean +/- standard error; $p<0.05$, **$p<0.01$, ***$p<0.001$; n=11-12 mice/group.

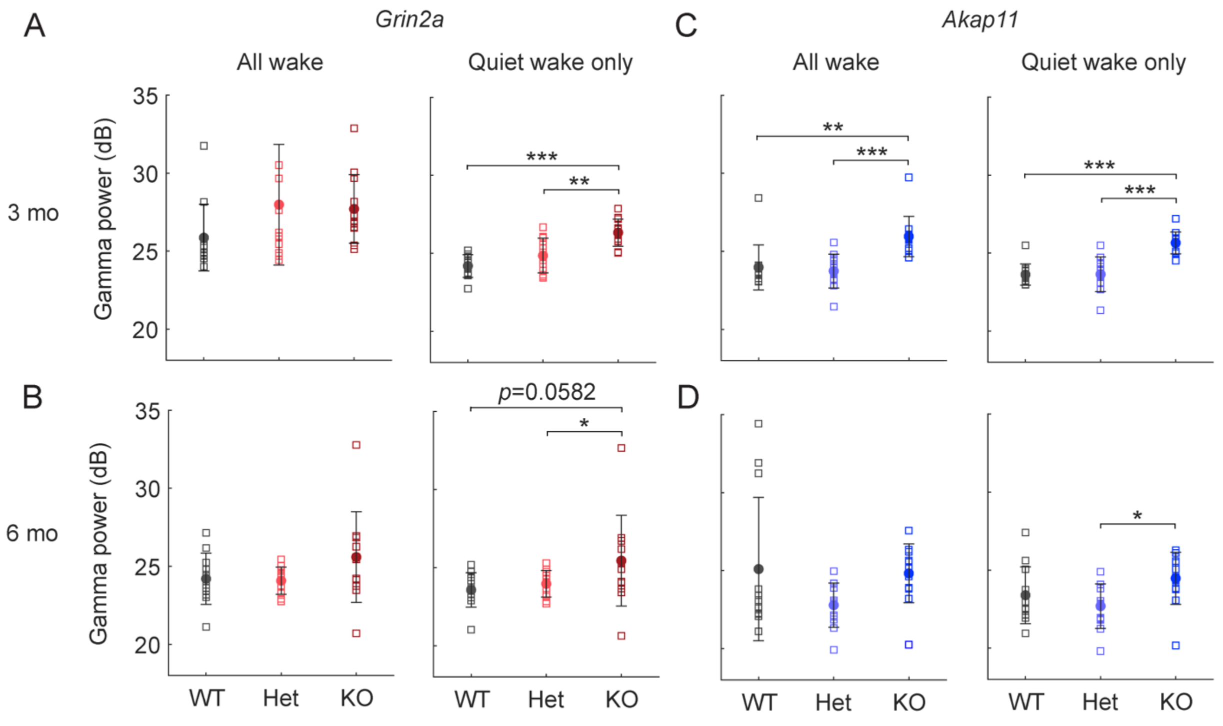


**Figure S5. Gamma oscillations in *Grin2a* and *Akap11* mutant mice during wake.** Absolute PSD for 3- and 6-month *Grin2a* (A-B) and *Akap11* (C-D) mutants computed for gamma (γ), 30-50 Hz, for wake states during the dark cycle. Results are shown for wake and quiet wake states (periods of presumed immobility during wakefulness; see Supplementary Methods for more details). During wake, *Akap11*$^{-/-}$ animals showed increased gamma power compared to WT littermates at 3 months (p=5.06e-04), with heterozygous animals similar to WT. Analysis of quiet wake states revealed additional increases in gamma power in *Grin2a*$^{-/-}$ animals at 3 months (p=6.87e-06) and a trend towards increased gamma in KO at 6 months (p=0.0582), similar to NREM (Figure 3) and REM sleep (Figure S3-4). Both *Grin2a* and *Akap11* heterozygous mutants exhibited similar gamma power as WT littermates. Error bars show mean +/- standard error; *p<0.05, **p<0.01, ***p<0.001; n=11-12 mice/group.

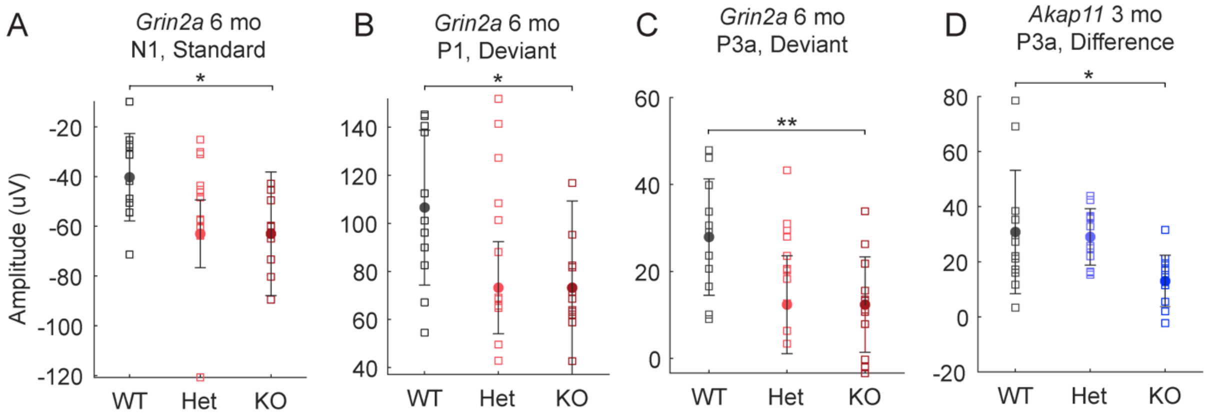


**Figure S6. Peak amplitudes during the mismatch negativity test in *Grin2a* and *Akap11* mutants.** Quantification of peak amplitudes for event-related potential components in the mismatch negativity test (Figure 6); significant results are shown for A) *Grin2a*, 6 months, N1 response to standard tones; B) *Grin2a*, 6 months, P1 and C) P3a response to deviant tones; D) *Akap11*, 3 months, P3a response in difference waveform [the response to deviant tones minus the response to standard tones. Error bars indicate +/- standard error; p<0.05, **p<0.01; n=11-12 mice/group.
